# Meta-analytic maps of brain activity evoked by cognitive function diverge from resting state networks

**DOI:** 10.1101/2021.05.05.442861

**Authors:** Matías Palmucci, Enzo Tagliazucchi

## Abstract

Spontaneous human brain activity is organized into resting state networks (RSN), complex patterns of synchronized activity that account for the majority of brain metabolism. The correspondence between these patterns and those elicited by task performance suggests that spontaneous brain activity originates from the stream of ongoing cognitive processing. Here we investigate a large number of meta-analytic activation maps obtained from Neurosynth (www.neurosynth.org) to establish that task-rest similarity can be inflated by two potential sources of bias. Applying a hierarchical module detection algorithm to a network representation of activation map similarity, we showed that the correspondence between RSN and task-evoked activity tends to hold only for the largest spatial scales. Second, we established that this correspondence is biased by the inclusion of maps related to neuroanatomical terms (e.g. “parietal”, “occipital”, “cingulate”, etc.). Our results challenge the cognitive origin of spontaneous brain activity, suggesting that anatomically-constrained homeostatic processes could also play an important role in the inception and shaping of human resting state activity fluctuations.

## Introduction

Brain metabolism consumes near 20% of the daily energy intake and this demand increases only 4% during cognitive effort, highlighting that most of the energetic cost of brain activity is related to the spontaneous baseline fluctuations seen during rest (i.e. in absence of sensory stimulation and instructions to perform cognitively effortful tasks) (Raichle, 2006). Due to this high metabolic cost it has been conjectured that spontaneous brain activity is evolutionary advantageous, playing a central role to support normal cerebral function (Bullmore and Sporns, 2012).

The field of resting state neuroimaging addresses this conjecture by providing a characterization of spontaneous brain activity in health and disease, and during different levels of arousal and responsiveness. The study of coordinated functional magnetic resonance imaging (fMRI) signals has evolved considerably since the pioneering study of Biswal and colleagues, who demonstrated significant spontaneous synchronization (i.e. functional connectivity) between anatomically-distant brain regions (Biswal et al., 1995). Data-driven methods such as independent component analysis (ICA) have been used to reveal large-scale patterns of synchronized spontaneous activity, which are frequently called resting state networks (RSN) (Beckmann et al., 2005). These networks outline well-known functional systems of the brain, such as the medial and lateral visual cortices, the auditory and sensorimotor cortices, and temporal-frontal-parietal networks linked to spontaneous conscious mentation (default mode network), executive control, and attention. These patterns can be reproduced robustly across species (Hutchison et al., 2011; Sforazzini et al., 2014) and individuals (Damoiseaux et al., 2006), can be revealed from single subject fMRI recordings (Hacker et al., 2013), present idiosyncratic alterations in several neurological and psychiatric conditions (Greicius, 2008), and are correlated with electrophysiological measurements of brain activity (Laufs et al., 2003; Mantini et al., 2007). Taken together, these results highlight the neurobiological significance of RSN and the relevance of understanding their origin and function.

It has been hypothesized that RSN originate from spontaneous cognition taking place during conscious wakefulness (also known as mind-wandering) (Gonzalez-Castillo et al., 2021). This hypothesis is compatible with the small increase in metabolism seen during effortful tasks, since spontaneous brain activity would already reflect ongoing cognitive processing; furthermore, it is also consistent with studies showing that resting state activity can be used to decode mental states (Richiardi et al., 2011; Gonzalez-Castillo et al., 2015), as well as to predict performance in motor and perceptual tasks (Boly et al., 2007; Hesselmann et al., 2008; Kannurpatti et al., 2012; Sadaghiani et al., 2015; Herszage et al., 2020). A seminal study published by Smith and colleagues demonstrated a direct correspondence between the spatial configuration of RSN and task-evoked activity patterns, obtained by applying ICA to a large database of brain activation maps (www.brainmap.org) (Smith et al., 2009). The authors were capable of matching all major RSN to the independent components extracted from this database, resulting in the data-driven assignment of a functional role to each network, thus supporting the hypothesis of RSN originating from spontaneous cognition.

The findings of Smith et al. have been corroborated using other methods and databases (Toro et al., 2008; Laird et al., 2013; Cole et al., 2014; Nickerson et al., 2018); in particular, using the Neurosynth platform (Di et al., 2013), which combines natural language processing and machine learning to mine activation coordinates from a very large set of published articles (>15.000 articles at the time the present analyses were conducted) (Yarkoni et al., 2011). The similarity between spontaneous and resting state activity patterns is generally accepted, having informed the influential theoretical perspective of a “dynamic baseline”, i.e. spontaneous brain activity as a trajectory unfolding in the proximity of attractors representing task-related brain configurations, thus being capable of rapidly transitioning towards these configurations as a consequence of environmental threats and demands (Deco et al., 2011). While attractive, this account is challenged by the persistence of RSN during unconscious states, such as deep sleep, general anesthesia, and unresponsive wakefulness syndrome in brain-injured patients (Boveroux et al., 2010; Soddu et al., 2011; Tagliazucchi et al., 2013). These observations are consistent with the alternative hypothesis of RSN originating from homeostatic processes constrained by brain anatomy, especially by long-range structural connections between cortical and sub-cortical regions (Greicius et al., 2009) which limit the disintegration of RSN during unresponsiveness (Barttfeld et al., 2015; Tagliazucchi et al., 2016).

Here we revisited the correspondence between evoked and spontaneous activity from a critical perspective, establishing that this correspondence could be inflated by two sources of bias which failed to be considered in most previous studies. Instead of focusing on task-rest similarity at the coarsest spatial resolution by default, we obtained a hierarchical clustering of the Neurosynth activation maps and showed that this similarity held only for the largest spatial scale. We also showed that this correspondence was biased by the inclusion of maps related to anatomical terms (e.g. “parietal”, “occipital”, “cingulate”, etc.). Taken together, these results challenge the cognitive origin of spontaneous brain activity, suggesting that anatomically-constrained homeostatic processes are also an important contributing factor.

## Materials and methods

### Neurosynth association test maps

Neurosynth is an automated framework to extract activation coordinates and synthesize association and uniformity test maps based on text-mining and machine learning techniques. Each of these maps is linked to a term that is frequently found in the neuroimaging literature, e.g. “language”, “parietal”, “motor”. The uniformity test map of a given term contains high z-score values in a brain region if the articles using that term consistently report activations in that region. The association test map of a given term is similar, except that it also considers the likelihood of the region being activated in articles that do not include the term, thus controlling for the base rate occurrence of the terms and increasing the specificity of the resulting activation maps. For more information, see www.neurosynth.org or the original publication by Yarkoni and colleagues (Yarkoni et al., 2011).

A total of 3072 association test maps were downloaded from Neurosynth, each corresponding to an individual term in the database. Repeated or redundant terms were discarded after manually verifying a high degree of similarity between the associated maps. The resulting terms were divided into a set that referenced cognitive processes (e.g. “vision”, “language”, “emotion”, “decision making”; N=400) and another that contained neuroanatomical terms (e.g. “occipital”, “parietal”, “temporal”, “pre-frontal”; N=409). Terms that did not fit into either category (e.g. names of neuropsychiatric diseases) were discarded from further analysis.

### Map similarity network

Two weighted undirected networks were constructed from the association test maps linked to cognitive and neuroanatomical terms, respectively. Edge weights reflected the degree of spatial similarity between pairs of maps. To estimate this similarity, each map was first thresholded using a false discovery rate (FDR) criterion of 0.01 (i.e. 1% false positives) and then downsampled from 2× 2 × 2 mm to 4× 4 × 4 mm. Given maps M_i_ and M_j_ (corresponding to the i-th and j-th nodes of the network), their similarity was computed using the Jaccard index, i.e. W_ij_ = |M_i_ ⋂ M_j_|/|M_i_ ∪ M_j_|, where W_ij_ represents the weighted adjacency matrix of the network.

### Hierarchical modular decomposition of map similarity networks

An algorithm based on the spectral decomposition of the weighted adjacency matrix (Newman, 2006) was employed to obtain modules at different scales. This algorithm explores partitions into modules with the purpose of maximizing the generalized definition of Newman’s modularity, Q = (1/2m) ∑_ij_(W_ij_ — γ k_i_k_j_/2m)δ_gigj_. In this equation, m is the total number of edges in the network, W_ij_ is the weighted adjacency matrix, k_i_ is the sum of weights attached to the i-th node, and δ_gigj_ = 1 if nodes i-th and j-th belong to the same module in the partition, and 0 otherwise. The parameter γ determines the resolution of the modular decomposition, i.e. the characteristic size of the resulting modules. By increasing the value of γ, the algorithm yields progressively smaller modules which are hierarchically contained into those obtained using smaller γ values. A diagram outlining this procedure is shown in Fig. 1.

**Figure 1.**
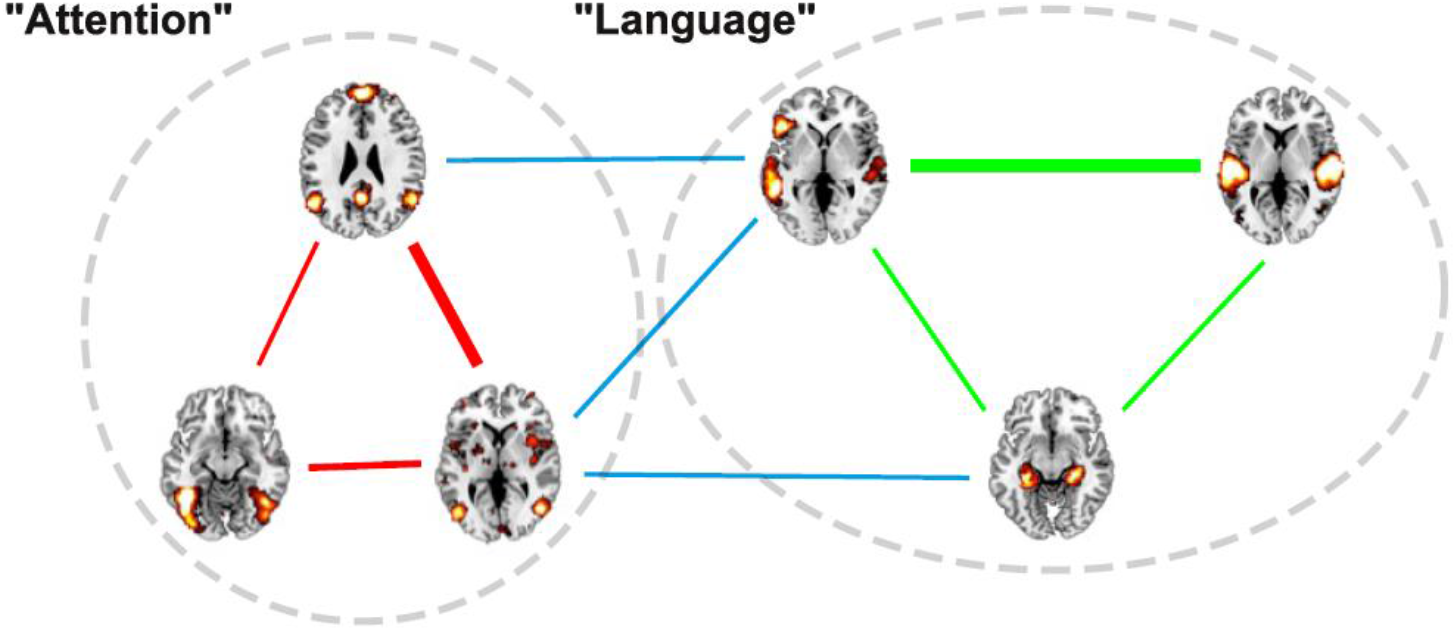
Schematic of the process followed to cluster the Neurosynth maps using a module detection algorithm. Each association map corresponds to a node in a network where the edge weights indicate the degree of spatial similarity (computed using Jaccard’s index). In this example, edge weights are represented by the width of the lines joining the maps. The network can be decomposed into two modules, one containing maps generally associated with “attention” (within-module links shown in red) and the other containing maps generally associated with “language” (within-module links in green). Between-module links are shown in light blue.

For each value of γ, the modularity maximization algorithm grouped nodes (maps) into disjoints sets according to their pairwise spatial similarity. Each module was summarized by computing the average of all maps it contained. Afterwards, the Pearson’s linear correlation coefficient between the average map and each of the maps was computed, and a word cloud representation of the associated terms was constructed using the correlation coefficient to determine the size of each term.

### Similarity to RSN

The average map obtained from each module was compared to the original set of 8 RSN published by Beckmann and colleagues (Beckmann et al., 2005). Average maps were characterized by its linear correlation with the following RSN: medial visual (VisM), medial lateral (VisL), auditory (Aud), sensorimotor (SM), default mode (DM), executive control (EC), dorsal right (DorR), dorsal left (DorL). A threshold of R = ±0.3 was used to determine a sufficiently large effect size.

## Results

The main results of our analysis are summarized in Fig. 2 (for the terms related to cognitive functions) and Fig. 3 (for the neuroanatomical terms). Each figure represents a hierarchical set of modules obtained by progressively increasing the resolution parameter γ. Each module is summarized by the average of all their activation maps, by the word cloud indicating the relative importance of their terms, and by the similarity (i.e. linear correlation coefficient) between their average map and the RSN defined by Beckmann and colleagues (Beckmann et al., 2005). As γ increases the modules can unfold into smaller sub-modules; in Figs. 2 and 3 this process is represented by a colored outline which contains both the “parent” and “children” modules.

**Figure 2.**
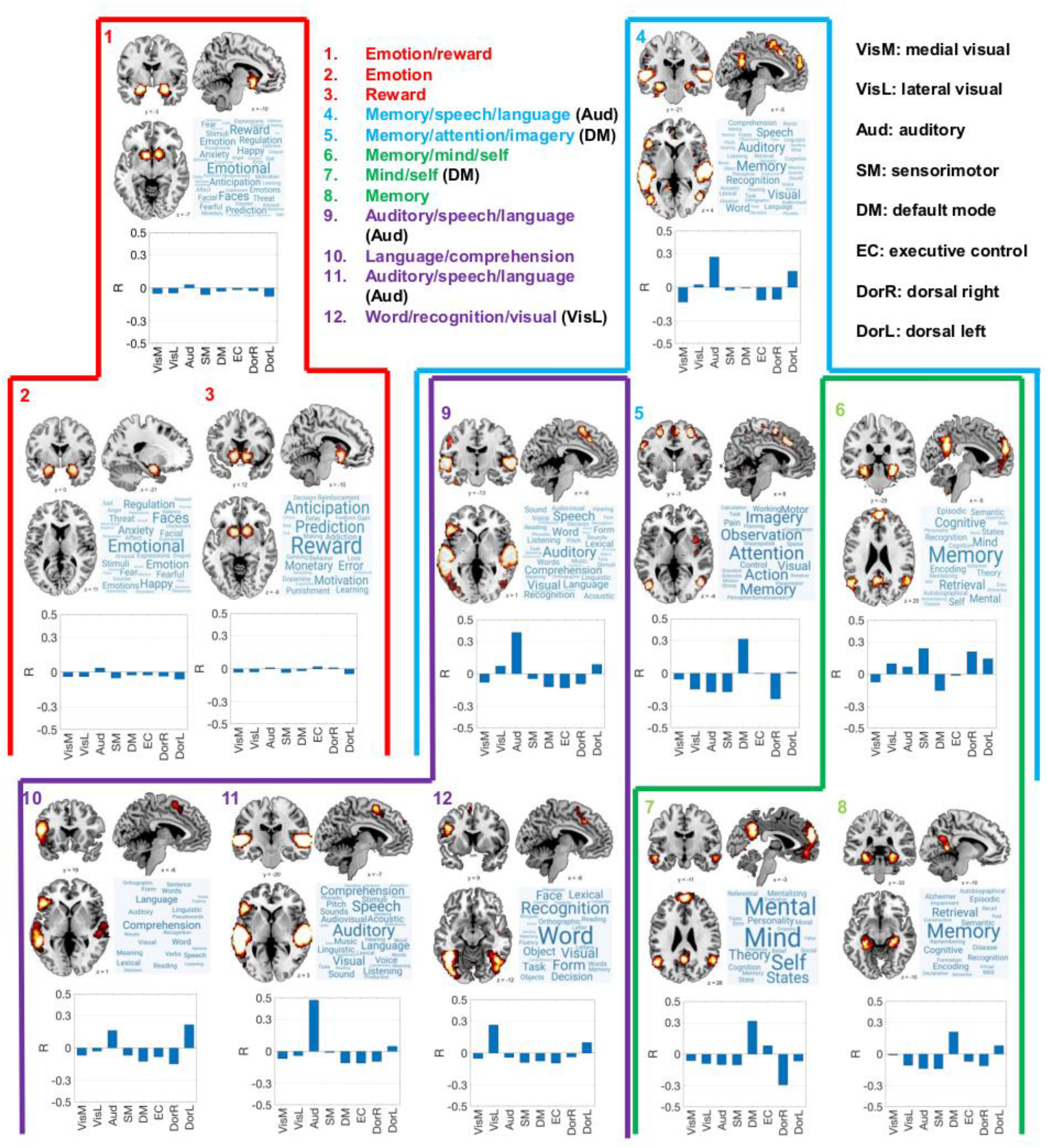
Hierarchical decomposition of Neurosynth association test maps corresponding to terms linked to cognitive functions. Each number indicates a module, represented by its average spatial map, its word cloud representation of term importance, and the correlation between the average map and the RSN published by Beckmann and colleagues. The hierarchical unfolding of the maps is represented by the colored outlines, i.e. map 1 unfolded into maps 2 and 3 as the resolution parameter γ was increased; map 4 into maps 9, 5 and 6; map 9 into maps 10, 11 and 12; map 6 into maps 7 and 8. Insets contain the name of each module as inferred from the word clouds. The abbreviated names of RSN presenting significant (|R| ≥ ±0.3) correlations with the average map appear next to the module name.

**Figure 3.**
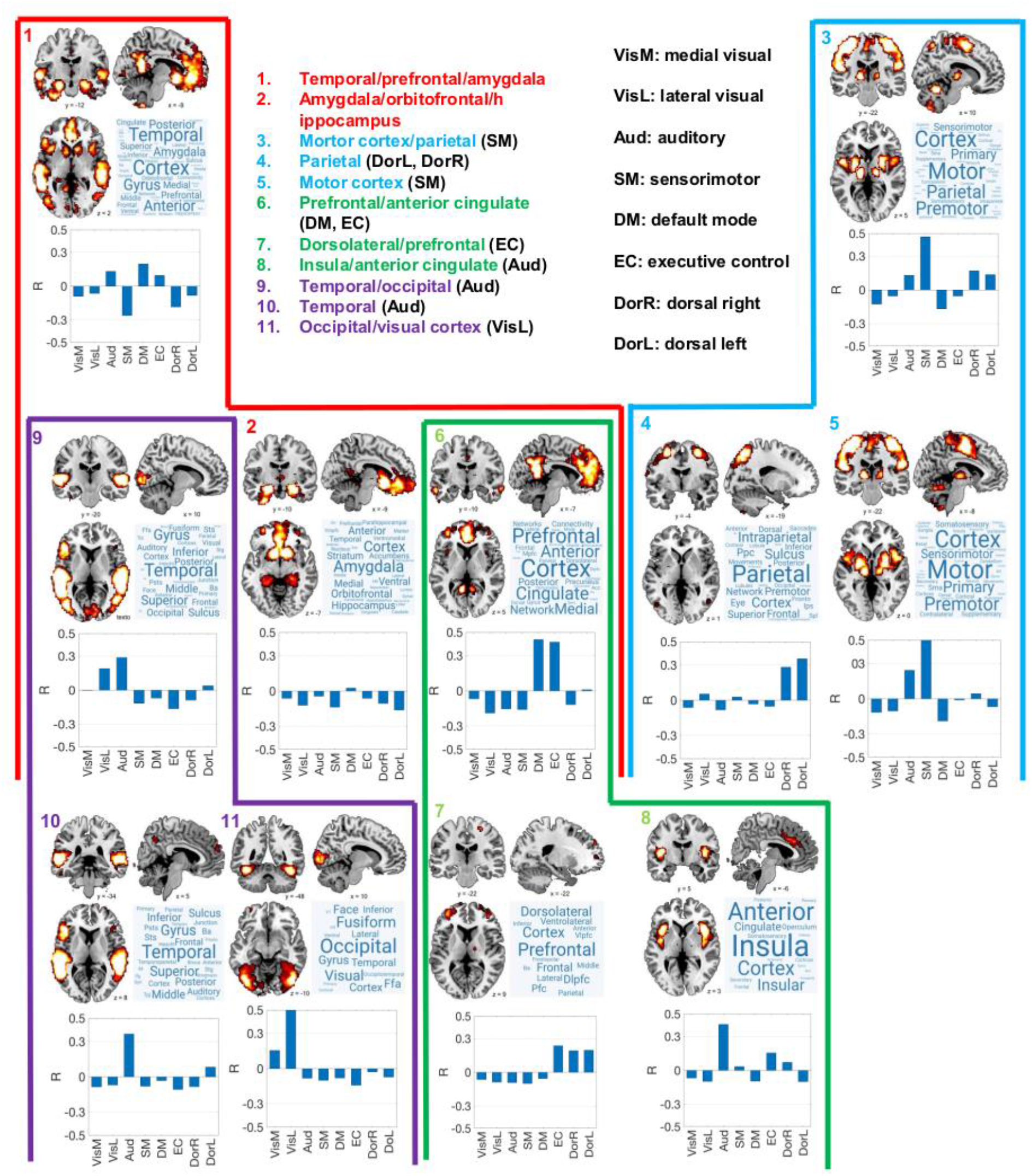
Hierarchical decomposition of Neurosynth association test maps corresponding to neuroanatomical terms only. Each number indicates a module, represented by its average spatial map, its word cloud representation of term importance, and the correlation between the average map and the RSN published by Beckmann and colleagues. The hierarchical unfolding of the maps is represented by the colored outlines, i.e. map 1 unfolded into maps 9, 2 and 6 as the resolution parameter γ was increased; map 3 into maps 4 and 5; map 9 into maps 10 and 11; map 6 into maps 7 and 8. Insets contain the name of each module as inferred from the word clouds. The abbreviated names of RSN presenting significant (|R| ≥ ±0.3) correlations with the average map appear next to the module name.

### Terms linked to cognitive functions

Association test maps corresponding to terms linked to cognitive functions were first divided into two modules: emotion/reward and memory/speech/language (Fig. 1, red hierarchy). The first of these two contains the amygdala and nucleus accumbens, and is related to terms such as “emotion”, “happy”, “anxiety”, “reward”, “anticipation” and “prediction”. As γ was increased, this module unfolded into one specific to emotion-related terms (map 2, including the amygdala) and another specific to reward-related terms (map 3, including the nucleus accumbens). None of the maps in this hierarchy presented significant similarity to any of the RSN.

The module related to memory/speech/language presented a significant overlap with the auditory RSN, but also included voxels within parietal, cingulate and hippocampal regions (Fig. 1, blue hierarchy). This module unfolded into three maps: one related to speech and language (map 9), another related to memory, attention and imagery (map 5) and a third map related to memory and self-processing (map 6). Map 9 presented a significant overlap with the auditory RSN and contained the temporal lobes, but excluded the bilateral hippocampus (hence the term “memory” is absent from its word cloud). Map 5 is related to cortical regions involved with memory, imagery and attention, and presents a significant overlap with the DMN. Map 6 contains the bilateral hippocampi and related regions, consistently with the words represented in its word cloud (“memory”, “encoding”, “retrieval”, “episodic”, etc.); however, this module did not present a significant correlation with any of the RSN.

Maps 9 and 6 could be further subdivided by increasing the resolution parameter γ (Fig. 2, purple and green hierarchies). The auditory/speech/language module (map 9) was divided into a module specific to language comprehension (map 10), another specific to auditory perception (map 11, presenting a high overlap with the auditory RSN) and a map linked to visual object recognition, including terms such as “visual”, “word”, “form”, “face”, “recognition”, “object”, and others (map 12, overlapping with the lateral visual RSN).

Finally, map 6 (memory/mind/self) unfolded into map 7 (cortical regions related to self-consciousness and to theory of mind, overlapping with the DMN) and map 8 (hippocampal and related regions, related to memory encoding and retrieval).

### Neuroanatomical terms

The network of activation maps linked to neuroanatomical terms (Fig. 3) was first divided into two modules, roughly representing temporal, frontal and pre-frontal and certain subcortical structures (map 1) and another representing motor and sensory areas in the prefrontal and parietal lobe, respectively (map 3, significant overlap with the sensorimotor RSN).

The first of these two modules (Fig. 3, red hierarchy) unfolded into a map outlining the temporal and occipital lobes (map 9, overlapping with the temporal RSN), another representing the amygdala/orbitofrontal cortex/hippocampus (map 2) and a third module including the prefrontal cortex and the anterior cingulate (overlapping with the default mode and executive control RSN).

The second of the two initial modules, the motor cortex/parietal module (Fig. 3, blue hierarchy), unfolded into a module including parietal regions (map 4, overlapping with left and right dorsal RSN) and another outlining motor and prefrontal areas (map 5, overlapping with the sensorimotor RSN).

Map 9 (Fig. 3, purple hierarchy) was further subdivided into a module specific to the temporal lobe (map 10, overlapping with the auditory RSN) and an occipital module including the fusiform gyrus (map 11, correlated with the lateral visual RSN). Map 6 (Fig. 3, green hierarchy) unfolded into a prefrontal/dorsolateral map overlapping with the executive control RSN (map 7) and another map containing the insula and the anterior cingulate cortex (map 8) which correlated with the auditory RSN.

In summary, all except two of the modules obtained from neuroanatomical terms presented significant overlap with at least one RSN, compared to half the modules obtained from terms linked to cognitive functions.

## Discussion

The origin and function of spontaneous brain activity remain two of the most outstanding unsolved problems in the field of human neuroimaging. Early studies attempted to dismiss RSN as artefacts caused by brain vasculature and physiological noise (Birn, 2012; Tak et al., 2015; Tong et al., 2015), however, their neural origin received ample support from multimodal imaging studies conducted in humans and animal models (Laufs et al., 2003; Mantini et al., 2007; Shmuel and Leopold, 2008; Schölvinck et al., 2010). The hypothesis of unconstrained cognitive processing as the source of coordinated spontaneous activity fluctuations gained terrain after a seminal study showing that RSN could be put in correspondence with the independent components obtained from a large number of meta-analytic task activation maps (Smith et al., 2005). This result received support from the study of task co-activation networks, as well as from studies which directly compared independent components obtained during task performance vs. those obtained during rest (Toro et al., 2008; Laird et al., 2013; Di et al., 2013; Cole et al., 2014; Nickerson et al., 2018)

The hypothesis of spontaneous cognitive processing as the origin of RSN is very attractive from a theoretical viewpoint. As a self-organized non-equilibrium dynamical system, the human brain continuously transverses different large-scale activity patterns (Cavanna et al., 2018); in turn, the proximity of these patterns to those seen during cognitive processing ensures rapid reactivity upon environmental threats and demands (Deco et al., 2011). This proximity can be realized by different dynamical mechanisms, for instance, attractor ruins that drive the configuration of brain activity towards patterns reflecting different activation maps (Deco and Jirsa, 2012). A problem with this hypothesis is that other factors seem to play an important role to shape resting state activity fluctuations, even during brain states characterized by diminished or absent conscious cognition. For instance, resting state activity fluctuations are constrained by large-scale anatomical connectivity (Greicius et al., 2009) and local gene expression profiles (Wang et al., 2015), which suggest the presence of background homeostatic processes unfolding independently of task-evoked activity.

We attempted to replicate the meta-analytic results published by Smith and colleagues using a different method applied to a newer database of brain activations (Yarkoni et al., 2011). The main distinctive features of our study comprised the independent analysis of maps related to cognitive vs. neuroanatomical terms, and the hierarchical grouping of overlapping activation maps. Our analysis revealed that patterns similar to RSN tend to appear as a consequence of maps linked to anatomical terms, and that the similarities driven by proper cognitive activation maps are scale-dependent, i.e. at finer scales, the meta-analytic maps reflect activations localized at specific neuroanatomical structures (e. g. amygdala, hippocampus) that diverge from the distributed nature of RSN.

We highlight that our analyses only undermine the correspondence between RSN and task-related maps derived from large meta-analytic databases. Other studies (including a replication of Smith et al., 2008, see Nickerson et al., 2018) found similarities between resting state activity and activity patterns elicited by experimental tasks in a group of participants, i.e. without resorting to the analysis of BrainMap or Neurosynth. A main limitation of these analyses is the restricted number of tasks that could be examined, complicating the direct comparison with our results, especially considering that some of these studies found discrepancies between resting state and evoked brain activity (Bolt et al., 2017).

The hierarchical organization of brain structure and function has been repeatedly demonstrated in humans and other animal models by means of an ample spectrum of experimental techniques (Bassett et al., 2008; Meunier et al., 2009; Smith et al., 2019; Xu et al., 2020). Our clustering of the Neurosynth database was consistent with this observation: by increasing the resolution parameter of the module detection algorithm, we were able to unfold larger modules (i.e. clusters) into sub-modules, which progressively reflected more specialized cognitive functions restricted to precise neuroanatomical boundaries. For example, a DMN-like map linked to self-consciousness, social cognition and memory was subdivided into a cortical fronto-parietal network of regions strongly resembling the DMN (bilateral precuneus, temporo-parietal junction and orbitofrontal cortex) linked to the first two aforementioned functions, and another network restricted to regions within the medial temporal lobe (i.e. hippocampus, parahippocampus). The task-positive network, comprising sensory, motor, executive and attentional regions, also subdivided as expected, with end clusters reflecting specific functions such as auditory perception, object recognition (secondary visual areas), and language production and comprehension. In contrast, hierarchical clustering of resting state brain activity yielded distributed spatial patterns which were very similar to RSN (or parts of RSN) and could not be easily put in correspondence with the more specific maps presented in Fig. 2 (Lee et al., 2012; Wang and Lee, 2013; Bellec, 2013).

In summary, we showed that the hierarchical grouping of Neurosynth activation maps associated with different cognitive functions departs from the RSN patterns that are robustly reproduced in humans. This observation does not eliminate the possibility of task-evoked activity patterns being similar to spontaneous activity fluctuations; however, this should be shown by experiments designed for this purpose, ideally gathering resting state and task-evoked activity from the same sample of participants. Both spontaneous cognition and baseline physiological processes constrained by neuroanatomy are likely to play a role in the origin of RSN; these contributions could be disentangled by factoring individual anatomical connectivity into the analyses, as well as by manipulating the level and content of unconstrained cognition.

## Notes

### Competing Interest Statement

The authors have declared no competing interest.

